# Differentiation protocol and maturation shape the axial identity and synaptic state in human iPSC-derived spinal motor neurons

**DOI:** 10.64898/2026.09.14.751550

**Authors:** Alba Sansa, Vladimir A Zhemkov, Monica Feole, Jaquelyn M Villalba, Shaughn Bell, Clive N Svendsen

**Affiliations:** Cedars-Sinai Board of Governors Regenerative Medicine Institute, Los Angeles, CA, USA

## Abstract

Human induced pluripotent stem cell (iPSC)-derived motor neurons are widely used for disease modeling. Stem cells can be differentiated into motor neurons by exposure to specific patterning morphogens, which can ultimately determine terminal cellular composition and developmental state. We compared two small molecule workflows for human iPSC maturation into motor neurons: a rapid direct protocol (diMN protocol) and an extended protocol (purMN protocol), with both protocols matched for 20 days in maturation medium. While the diMN protocol generated mixed neural cultures, the purMN protocol provided cultures with greater motor neuron enrichment, stronger caudal spinal identity, broader synaptic and cholinergic programs, and reduced progenitor-associated signatures. By extending the purMN protocol to 32 days in maturation media, cultures developed reinforced spinal, postsynaptic, presynaptic, calcium-signaling, and cholinergic features. Across four independent reference frameworks, deconvolution aligned diMN cultures with anterior and progenitor-associated states and purMN cultures with spinal and post-mitotic states. In a microfluidic co-culture system, purMNs formed denser, more highly branched distal neurite networks and produced more innervated acetylcholine receptor clusters compared to diMNs. Revealing how purity and maturation jointly define the transcriptomic and structural state of human iPSC-derived spinal motor neuron cultures will help the field choose optimal models for studies of neurodevelopment and neurodegenerative diseases.

## Background

Human induced pluripotent stem cells (iPSCs) can be differentiated *in vitro* into cells of the central nervous system, providing experimental access to otherwise poorly accessible human cell types^1^. Specifically, human iPSC-derived spinal and cortical motor neurons (MNs) are widely used to model motor neuron development, injury and neurological diseases including Spinal Muscular Atrophy (SMA), Amyotrophic Lateral Sclerosis (ALS), and related neuromuscular disorders^2–10^. In terms of spinal MN differentiation strategies, most utilize developmental principles of neural tube patterning, through exposure to synthetic morphogens mimicking MN development by using dual-SMAD inhibition followed by caudalization with retinoic acid (RA) and ventralization through sonic hedgehog (SHH) signaling, thereby directing cells toward OLIG2-positive MN progenitor and post-mitotic spinal MN states^2^. A major advance in the field was the development of efficient methods to generate and expand highly pure spinal MN progenitors from iPSCs, which provided a reproducible framework for obtaining lineage-restricted MN cultures at-scale^4^.

Various differentiation protocols have evolved to achieve different practical goals. Some workflows prioritized high-purity MN cultures and tighter lineage specification by expanding patterned motor neuron progenitors or by optimizing small-molecule neuralization, caudalization, and ventralization steps. These high-purity protocols enriched for OLIG2-positive progenitors and postmitotic MNX1/HB9-, ISL1, and ChAT-positive cholinergic spinal MNs, often exceeding 70% MN identity, thereby decreasing signal-to-noise for MN-intrinsic programs^2,11,12^. Alternative more rapid, direct workflows were optimized for scalability and throughput, which yielded mixed cultures^3,6,13^ in which MNs represent a minority population alongside neural progenitors and other neural cell types^3,6,13^, potentially diluting MN-specific transcriptional signatures. This direct protocol has been particularly influential in consortium-led large-scale MN differentiation and disease modeling efforts across large numbers of patient-derived lines^3,6,13^.

While both high-purity and mixed-culture protocols are built on the same core developmental principles, the derived MN pools substantially vary between the protocol types. Here we rigorously benchmark side-by-side the biological outputs of two published human spinal MN differentiation strategies applied to the same human iPSC lines: an extended protocol for high-purity motor neuron (purMN) cultures^2,7,8^ and a rapid protocol for direct motor neuron (diMN) cultures that yields a more mixed neuronal population^3,6,13^. Immunocytochemistry and bulk RNA sequencing demonstrated that protocol choice and maturation time points influence MN identity, progenitor program suppression, rostro–caudal patterning, and the emergence of neuronal, synaptic, and metabolic transcriptional signatures. We show that high-purity differentiation combined with extended maturation is required to robustly engage neuronal and synaptic transcriptional programs, which are weak or absent in cultures where the proportion of MNs is lower. Clearly, differences in progenitor handling, purification strategy, maturation state, and final culture composition can all influence the transcriptomic identity of the resulting cells and, by extension, the development and interpretation of downstream phenotypes. The critical role that protocol selection and maturation duration play to obtain mature motor neuron states is crucial knowledge for studies to faithfully model motor neuron development, injury and disease.

## Materials and methods

### Differentiation of human induced pluripotent stem cells (iPSCs) into spinal cord MNs

Human peripheral blood mononuclear cell (PBMC)-derived iPSC lines were obtained from the Cedars-Sinai Biomanufacturing Center. Three control lines, CS8VTRiCTR-nxx (parent GUID: NEUXC258VTR), CS0YX7iCTR-nxx (parent GUID: NEUVZ050YX7), and EDi022-A (RRID: CVCL_VT08), were used for both diMN and purMN differentiation. Donor characteristics and available line-specific quality-control information, including reprogramming method, STR authentication, mycoplasma testing, karyotype, and pluripotency assessments, are provided in Table S1 and the corresponding repository records. The iPSC colonies were maintained on Matrigel-coated 10 cm dishes in mTeSR (StemCell Technologies) for three weeks according to standard Cedars-Sinai protocols. iPSCs were passaged every 4–5 days using Versene Solution (Gibco) at a split ratio of 1:6. Differentiations were initiated from working-bank stocks at passages 30-35, and cells underwent at least 3 passages after thawing before differentiation. Stocks were cryopreserved in CryoStor® CS10 (StemCell Technologies) and thawed onto Matrigel-coated dishes in mTeSR.

Once iPSC colonies reached 70-80% confluence (about 4-5 days after passaging), iPSCs were dissociated with Accutase (Gibco) and plated for differentiation into spinal motor neurons using two separate small-molecule workflows that both use early SMAD-pathway inhibition and WNT activation, but differ in the BMP inhibitor used during neural induction and differ in subsequent motor neuron progenitor expansion (Fig. 1a)^2,6–8,13^. The short direct MN (diMN) workflow involved a 12-day differentiation stage, using LDN193189/SB431542-based dual SMAD inhibition along with CHIR99021 during early induction, followed by retinoic acid (RA)/SAG-mediated caudal-ventral patterning. In contrast, the extended pure MN (purMN) workflow involved a 24-day differentiation stage, using DMH1/SB431542/CHIR99021 during neuroepithelial induction, followed by RA/purmorphamine/valproic acid-driven motor neuron progenitor expansion before terminal maturation. In both protocols, cells were maintained at 37°C with 5% CO_2_. Cultures were routinely monitored for contamination; dedicated bacteriostasis and fungistasis testing was not performed.

**Figure 1.**
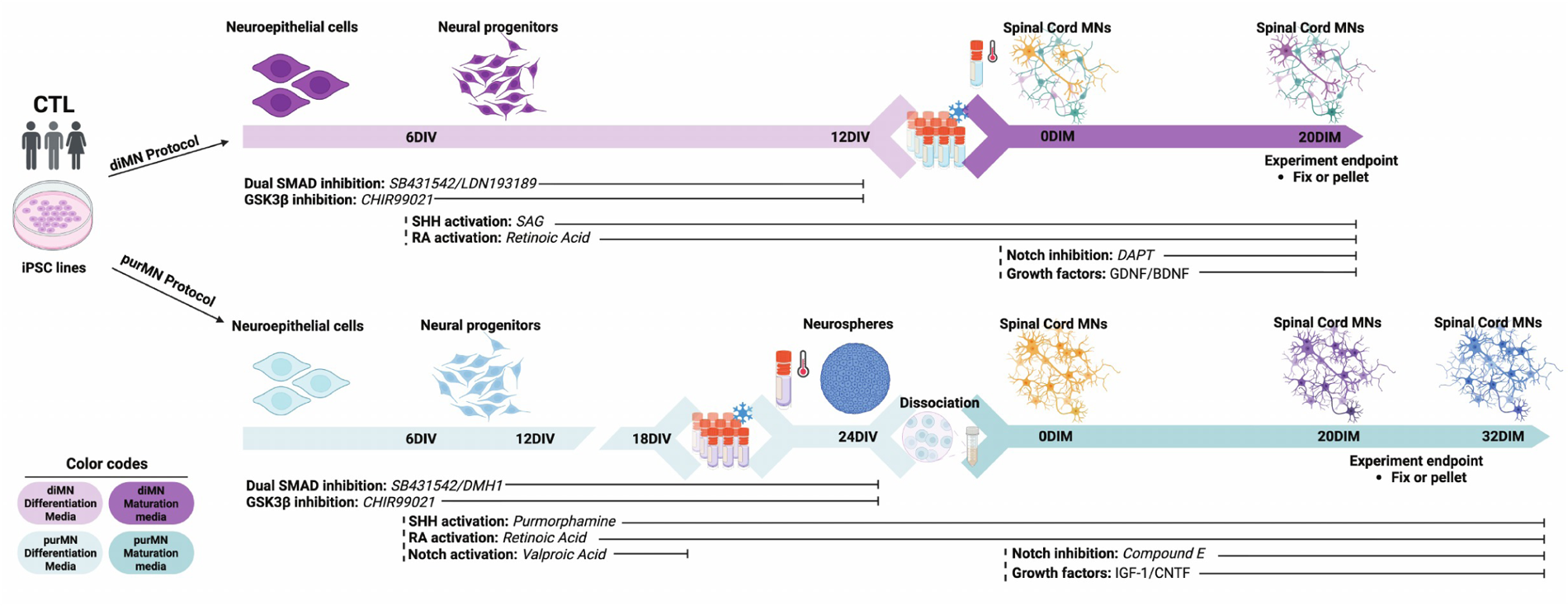
Schematic overview of the diMN and purMN differentiation and maturation workflows. Control human iPSC lines were differentiated using the diMN protocol (top) or purMN protocol (bottom). The diMN workflow generated neuroepithelial cells and neural progenitors over 12 days *in vitro* (DIV) using dual-SMAD inhibition, GSK3β inhibition, retinoic acid (RA)-mediated caudalization, and sonic hedgehog (SHH) pathway activation. Neural progenitors were cryopreserved at 12 days *in vitro* (DIV), subsequently thawed and seeded in diMN maturation medium containing DAPT, GDNF, and BDNF, and maintained for 20 days in maturation medium (DIM). In the purMN workflow, neuroepithelial cells and neural progenitors were generated using a distinct combination and timing of dual-SMAD inhibition, GSK3β inhibition, retinoic acid, SHH pathway activation, and valproic acid. Progenitors were cryopreserved at 18 DIV, subsequently thawed and maintained as neurospheres until 24 DIV, and then dissociated and seeded in purMN maturation medium containing Compound E, IGF-1, and CNTF. purMN cultures were maintained for either 20 or 32 DIM. Samples were fixed or collected as cell pellets at the indicated experimental endpoints. Colored bars distinguish the differentiation and maturation media used for each protocol.

#### diMN Protocol

Cells were differentiated to MNs as described^3,6,13^. Dissociated iPSCs were plated in Stage 1 media consisting of a base media (47.5% IMDM (Gibco), 47.5% F12 (Gibco), 1% NEAA (Gibco), 1% Pen/Strep (Gibco), 2% B27 (Gibco), 1% N2 (Gibco)) with the addition of 0.2 μM LDN193189 (Stemgent), 10 μM SB431542 (StemCell Technologies), and 3 μM CHIR99021 (Sigma Aldrich) and exchanged daily until day 6, when cells were incubated in Accutase (Gibco) for 5 minutes at 37°C, collected, centrifuged and plated in Stage 2 media consisting of Stage 1 media plus 0.1 μM all-trans RA (Sigma Aldrich) and 1 μM SAG (Cayman Chemicals). Media was exchanged daily until day 12, when cells were cryopreserved CryoStor® CS10 (StemCell Technologies).

For experiments, cells were thawed and plated in diMN maturation media consisting of the base media with the addition of 0.5 μM all-trans RA (Sigma Aldrich), 0.1 μM SAG (Cayman Chemicals), 0.1 μM Compound E (Millipore), 2.5 μM DAPT (Sigma Aldrich), 0.1 μM db-cAMP (Millipore), 200 ng/mL Ascorbic Acid (Sigma Aldrich), 10 ng/mL BDNF (PeproTech), 10 ng/mL GDNF (PeproTech). The diMN maturation media was exchanged every 3 days for 20 days, for a total of 32 days in culture.

#### purMN Protocol

Cells were differentiated to MNs as described ^2,7,8^. Dissociated iPSCs were plated in neuroepithelial induction medium (NEPIM): DM/F12:NBMPlus 1:1 supplemented with B27, Glutamax, and NEAA (all from Gibco), with the addition of 0.1 mM ascorbic acid (Sigma), 3μM CHIR99021, 2 μM SB431542 and 2 μM DMH1 (all from Cayman Chemicals) to generate neuroepithelial cells. After six days *in vitro*, neuroepithelial cells were dissociated and expanded in MN induction medium: NEPIM, plus 0.1 μM all-trans RA (Sigma), 0.5 μM purmorphamine (Cayman Chemicals) and 0.5 mM valproic acid (Sigma) was also added. Cells were cultured until day 18 to produce MN progenitors (MNPs), which were then cryopreserved. After thawing, MNPs formed free-floating neurospheres and were cultured in MN induction medium for 6 days. Neurospheres were then dissociated and plated on laminin-coated dishes in MN maturation medium: NBMPlus supplemented with B27, Glutamax, and NEAA (all from Gibco); 0.1 mM ascorbic acid (Sigma); 0.1 μM Compound E (Sigma), 1x CultureOne^TM^ (Gibco), 20 ng/ml CNTF (Peprotech), and 20 ng/ml IGF-1 (Peprotech) to induce MN maturation. The purMN maturation media was exchanged every 3 days for either 20 days or 32 days in culture.

### Skeletal muscle progenitor cells

One human iPSC line (CSC640iCTR-n6) was differentiated into skeletal muscle progenitor cells (SMPCs) with a directed skeletal myogenic differentiation workflow adapted from Shelton et al. and Hicks et al^14–16^. Briefly, iPSCs were dissociated to single cells and plated on Matrigel in mTeSR supplemented with 10 *μ*M HA-1077. Cells were then induced toward mesoderm in E6 medium containing 10 *μ*M CHIR99021, followed by sequential PAX3 and PAX7 myogenic specification stages. Cultures were maintained through PAX7 differentiation in DMEM/F12-based medium supplemented with N2, insulin-transferrin-selenium, IGF-1, and SB-431542.

Beginning at day 42, SMPCs were enriched by fluorescence-activated cell sorting (FACS) using the Hicks et al. surface-marker strategy^15^. Cultures were dissociated with TrypLE and collagenase, filtered, resuspended in FACS buffer, blocked with human Fc receptor block, and stained with antibodies against ERBB3/HER3, NGFR/CD271, and HNK1 together with a live/dead viability dye. Live ERBB3-positive, NGFR-positive, HNK1-negative cells were collected as SMPCs. Sorted SMPCs were expanded in SMPC expansion medium and maintained below 90% confluence before dissociation, counting, and seeding into the central compartment of NMJ-chip devices. Myogenic potential was demonstrated by differentiation into myosin heavy chain-positive myotubes in the NMJ-chip experiments.

### Assembly and culturing of the NMJ-chips

A three-compartment Polydimethylsiloxane (PDMS) Innsbruck silicon microfluidic system (Xona Microfluidics) mounted on glass coverslips was used to generate a neuromuscular junction (NMJ)-chip. The rectangular glass coverslips were cleaned in 1N HCl, washed in ddH2O and subsequently in 70% and 96% ethanol. Briefly, Xona chips were placed on top of glass coverslips, cured at 60°C for 3 hours, and sterilized by plasma-treatment, achieving efficient attachment of the chips onto the glass coverslips. Then, they were coated overnight with 0.1% poly-D-lysine, washed three times with ddH_2_O and left overnight at 37°C in ddH_2_O. Lateral compartments were coated with 20 *μ*g/mL laminin for at least 1h at 37°C and used for MN seeding. Either diMNs or purMNs were seeded at 50,000 cells per each lateral compartment in their respective maturation media but changing the basal media to BrainPhys media (StemCell Technologies). Once MN projections reached and expanded in the central compartment (typically within 2 weeks post-plating), the central compartment was coated with 4x Matrigel for 1 h at 37°C and then washed with BrainPhys media. Media was completely aspirated from the central compartment and 500,000 SMPC + 50,000 commercially available primary human dermal fibroblasts (Thermo Fisher, C0045C) were resuspended in 1 μL of myotube media (BrainPhys media, 1.2% N2, 0.5% B-27, 1x GlutaMAX, 20 ng/ml IGF-1, and 20 ng/mL CNTF). Primary human fibroblasts from a healthy individual were included to support SMPC adhesion (1:10 ratio SMPC:fibroblasts). The cell suspension was seeded into each the top and the bottom of the central channel openings. Cells were allowed to attach for 10 minutes at 37°C and then the central compartment was filled with myotube media. Half-media changes in each compartment were performed on alternating days. MNs and SMPCs were co-cultured for 10-14 days and fixed for immunocytochemistry.

### Transcriptomic analysis

The diMNs and purMNs were plated at 600,000 cells/well in laminin-coated six-well tissue-culture dishes (Corning Incorporated) for RNA-seq. Cultures were incubated at 37°C with Accutase (Gibco) for 5-10 minutes and diluted with an equal volume of the complete culture media in which they were grown. After pelleting cells at 200 × g for 5 min at 4°C, cells were resuspended in PBS and pelleted again. Total RNA was isolated using a RNeasy Micro Kit (Qiagen, Cat. No: 4004). RNA integrity was assessed using an Agilent 2100 Bioanalyzer, and only samples with RIN > 8 were selected for cDNA library construction. The SMART-Seq V4 Ultra Low RNA Input Kit for Sequencing (Takara Bio USA) was used for reverse transcription and generation of double-stranded cDNA for subsequent library preparation using the Nextera XT Library Preparation kit (Illumina). An input of 10 ng RNA was used for oligo(dT)-primed reverse transcription, followed by cDNA amplification and cleanup. Quantification of cDNA was performed using Qubit (Thermo Fisher Scientific). cDNA normalized to 80 pg/μl was fragmented, and sequencing primers were added simultaneously. A limiting-cycle PCR added Index 1 (i7) and Index 2 (i5) adapters, as well as the sequences required for cluster formation on the sequencing flow cell. Indexed libraries were pooled and cleaned, and the pooled library size was verified using an Agilent 2100 Bioanalyzer and quantified by Qubit. Libraries were sequenced on an Illumina NovaSeq 6000 to a depth of more than 30 million reads per sample. The same SMART-Seq V4/Nextera XT library-preparation workflow was used for all samples; sample-specific read layouts are reported in the GEO metadata. For the day-20 libraries, the FASTQ files supplied by the sequencing core were labeled as trimmed and represent the earliest available sequencing files; untrimmed instrument FASTQs were not available to the authors. Transcript-level quantification was performed with Salmon (v1.4.0) using a decoy-aware index based on the GRCh38 human transcriptome (GENCODE v38). Transcript estimates were imported into R with tximeta and summarized to the gene level. Downstream visualization and comparative analyses were performed on gene-level expression tables, including TPM-derived log2(TPM + 1) matrices used for principal component analysis, marker-gene visualization, enrichment analysis, and reference-guided signature scoring.

### Immunocytochemistry

The diMNs and purMNs were plated at 100,000 cells/well on 1cm^2^ laminin-coated glass coverslips placed into twenty-four-well dishes for immunofluorescence analysis. Cultured cells were fixed with 4% paraformaldehyde (Sigma) for 30 min. Cells were permeabilized with 0.5% Triton X-100 and incubated for 30 min with blocking solution (0.5% Triton X-100,10% BSA in PBS). Cells were incubated with primary antibodies (Table S2), followed by species-appropriate Alexa Fluor– conjugated secondary antibodies (Thermo Fisher Scientific). Nuclei were counterstained with 4′,6-diamidino-2-phenylindole (DAPI 1:10,000; Thermo Fisher Scientific). Coverslips were mounted on Superfrost glass slides using Fluoromount-G mounting medium (Southern Biotech) and allowed to cure overnight prior to imaging.

### Cell type quantification

Briefly, DAPI-stained nuclei were identified using automated thresholding and watershed segmentation to separate closely apposed nuclei. Total cell number was defined as the number of DAPI-positive nuclei per field. Astroglial markers (GFAP and S100B) were quantified at the field level rather than assigned to individual nuclei. For each image, marker-positive signal was identified using intensity thresholding, and the proportion of the image field occupied by GFAP- or S100p-positive signal was calculated relative to the total image area. Motor neuron markers (MNX1/Hb9, ISL1/2, or ChAT) were calculated by applying intensity thresholds to the corresponding fluorescence channels and scoring nuclei that overlapped with marker-positive signal.

Confocal images were acquired on a Nikon A1R microscope using an immersion oil 60× objective lense/1.4 NA Plan Apochromat. Cell population counts were performed using Fiji (ImageJ) with a custom macro to enable automated and unbiased analysis. For each condition, multiple non-overlapping fields were imaged per coverslip and across independent differentiations. Automated segmentation was visually inspected for accuracy, and threshold parameters were kept constant across all conditions within an experiment. Quantifications were pooled across fields and averaged per biological replicate prior to statistical analysis. All image analysis was performed blinded to experimental condition.

### Axonal morphology analysis in NMJ-chip

Distal axonal morphology in microfluidic NMJ-chips was quantified in Fiji from neurofilament heavy-stained images using a fixed rectangular region of interest (ROI) placed at the same position relative to the microgroove exits in all samples. Images were background-corrected, thresholded to generate binary axon masks, and cleaned with binary open/close operations using identical settings within each experiment. For neurite density and branching analysis, masks were skeletonized and analyzed with the Analyze Skeleton plugin to obtain total branch length, endpoints, and junctions. Axon density was calculated as total branch length per ROI area, and endpoint and junction densities were normalized to ROI area. For axonal thickness analysis, the same binary masks were processed with the Local Thickness plugin in 2D mode, and foreground thickness distributions were summarized per ROI as the weighted median thickness and weighted 90th percentile thickness. Each ROI was treated as one technical subsample, and all segmentations were visually checked by overlaying masks and skeletons onto the source images.

### Neurite and NMJ innervation analysis in NMJ-chip

Confocal images from microfluidic neuron-muscle co-cultures were analyzed in Fiji/ImageJ using a binary-mask intersection workflow to quantify alpha-bungarotoxin-labeled acetylcholine receptor (AChR) clusters contacted by motor neuron neurites. Multichannel images were opened in Fiji and, when necessary, split into separate AChR and neurite channels. All image (ROIs) within a given comparison were processed using identical analysis settings.

For AChR cluster segmentation, the alpha-bungarotoxin channel was background-corrected using Subtract Background with a rolling-ball radius of 20-50 pixels, thresholded using an automated method such as Otsu or Triangle, and converted to a binary mask. Masks were cleaned using Fill Holes and Open operations, and Watershed was applied when needed to separate touching clusters. Total AChR clusters were quantified using Analyze Particles with a minimum area threshold of 5 μm^2^ and a circularity range of 0.2-1.0. For neurite segmentation, the neurite channel was background-corrected using a rolling-ball radius of 50-80 pixels, enhanced with the Tubeness plugin, thresholded to a binary mask, and cleaned using Open and Close operations. The neurite mask was then dilated 3 times to compensate for small registration errors, synaptic cleft spacing, and z-projection misalignment.

Innervated AChR clusters were identified by logical intersection of the AChR mask and the dilated neurite mask using the Image Calculator (AND operation). The resulting contact mask was analyzed with Analyze Particles using the same size threshold applied to the total AChR mask to obtain the number of innervated clusters per ROI. Percent neuromuscular innervation was calculated as innervated clusters divided by total AChR clusters multiplied by 100. Where required, contact area was measured directly from the AChR-neurite intersection mask and normalized to individual AChR-cluster area using ROI-based measurements. Segmentation accuracy was verified visually by overlaying masks onto the source images to minimize false-positive assignments.

### Statistical analysis

Statistical analyses were performed using GraphPad Prism and R, with *p* < 0.05 considered statistically significant. Transcriptomic pairwise gene-level comparisons were assessed with limma on log2(TPM + 1) values followed by Benjamini-Hochberg correction; genes were considered significantly differentially expressed when BH-adjusted *p* < 0.05 and |logFC| >= 1. Pairwise significance in targeted synaptic, astrocyte, and related marker-family panels was displayed only when both the adjusted p-value threshold and the predefined effect-size threshold were met. Imaging, neurite, and AChR-cluster analyses used the statistical tests specified in the figure legends. Error bars represent standard error of the mean (SEM).

### Paper-guided deconvolution and cross-reference analysis

Bulk RNA-seq profiles were compared with four published single-cell reference frameworks, grouped into two categories. The developing human spinal-cord datasets from Rayon et al.^17^ and Andersen et al.^18^ were used as *in vivo* developmental benchmarks. The Ho et al.^19^ and Lall et al.^20^ datasets were analyzed separately as *in vitro* contextual references because their cellular states reflect specific differentiation and culture conditions. Ho et al. used a rapid conventional-culture differentiation that generated mixed cranial and spinal neuronal and progenitor states, whereas Lall et al. used diMN-derived progenitors matured under flow in a microfluidic spinal-cord chip with endothelial-like cells. Consequently, scores derived from Ho and Lall were interpreted as protocol- and culture-context alignments.

For each framework, cell-state and regional marker sets were curated from the published annotations and reported marker genes. Gene-level TPM values were log2(TPM + 1)-transformed and standardized across samples for each gene. A score was calculated for each state and sample as the mean standardized expression of its positive marker genes, with subtraction of the mean negative-marker score where applicable. Scores were converted to relative weights by applying a softmax transformation separately within each reference panel and sample, such that the weights summed to one. Group-level weights were calculated across three control iPSC lines per condition. These values represent relative alignment of the bulk transcriptomes with published marker programs and should not be interpreted as direct estimates of cell fractions.

Group differences for each state were assessed by one-way ANOVA, followed by Benjamini-Hochberg correction across states within each reference panel. Pairwise group comparisons were performed using Welch’s two-sample t-tests, with Benjamini-Hochberg correction across the three contrasts within each state. Kruskal-Wallis tests, with corresponding Benjamini-Hochberg correction across states, were performed in parallel as a nonparametric sensitivity analysis.

### Figure design and visualization

Graphical schematics were prepared with BioRender and Adobe Illustrator. Transcriptomic plots, enrichment figures, HOX/axial identity panels, synaptic and astrocyte marker panels, and deconvolution visualizations were generated in R and Codex, and assembled in Adobe Illustrator.

## Results

### High-purity protocol yields purMN cultures show a more spinal and less progenitor-like transcriptome than diMN cultures

To benchmark how protocol choice shapes culture composition, three control human iPSC lines were differentiated following either the diMN protocol or the purMN protocol (**Figure 1**). We first used bulk RNA-seq to compare the transcriptomes from diMN cultures and purMN cultures following the same 20 days in maturation medium (DIM) (**Figure 2a**). Principal-component analysis separated the two conditions (**Figure 2b**), and differential expression analysis showed that diMNs retained stronger progenitor and early neural-associated genes, whereas purMNs upregulated multiple caudal HOX genes and neuronal specialization-associated transcripts (**Figure 2c**). This divergence was also evident at the level of axial identity. The HOX heatmap showed that diMNs retained a more anterior/cervical-weighted pattern, whereas purMNs shifted toward a more caudal spinal profile with stronger brachial/thoracic-associated HOX expression (**Figure 2d**). Tissue-enrichment analysis linked diMNs to embryonic and neuroepithelial contexts (**Figure 2e**), and purMNs to spinal cord, MN, and other differentiated nervous system signatures (**Figure 2f**).

**Figure 2.**
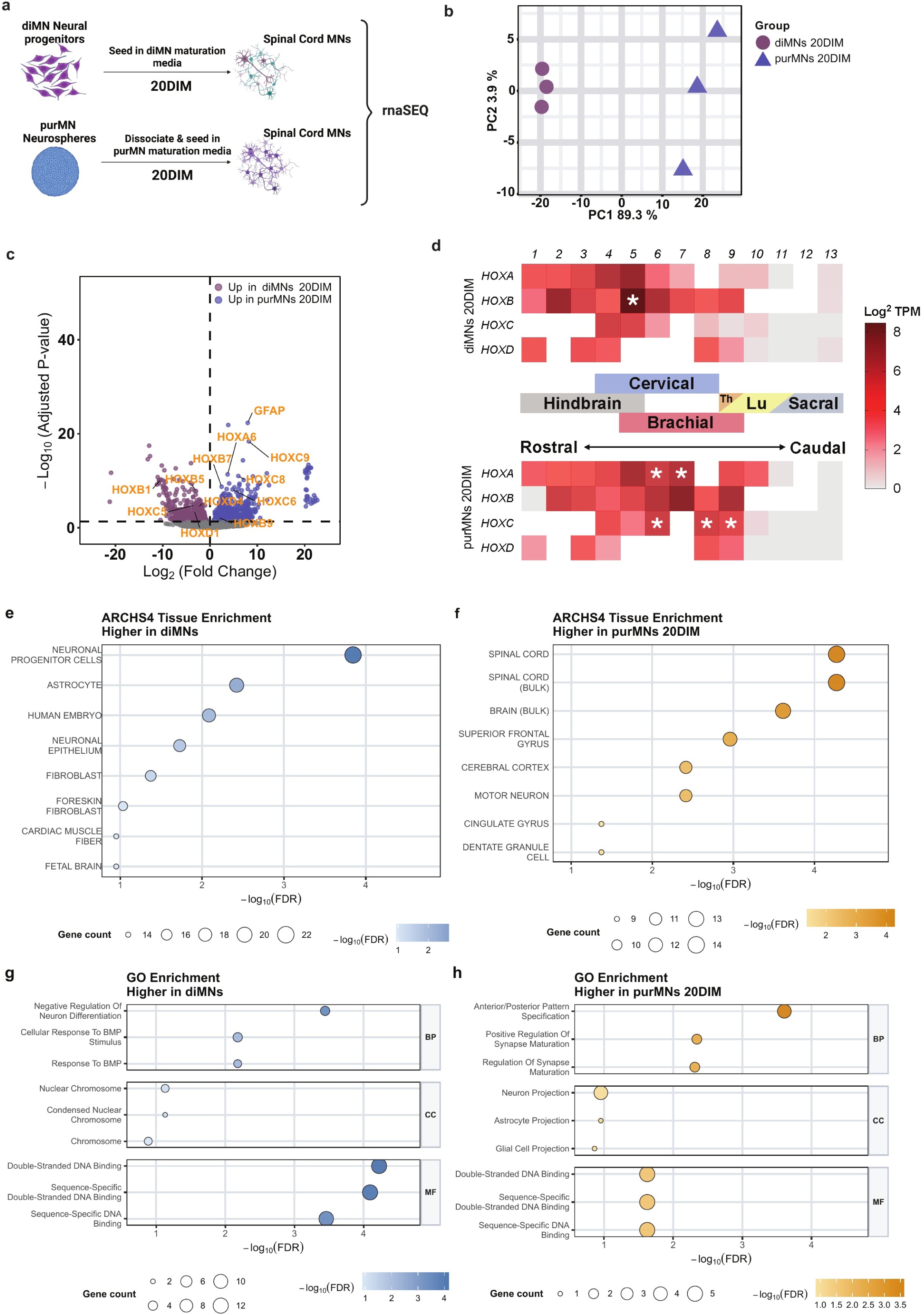
High-purity spinal motor neuron differentiation suppresses non-motor-neuron programs and strengthens caudal identity at matched 20DIM. (a) Experimental design for the two-group RNA-seq comparison between diMNs 20DIM and purMNs 20DIM. (b) Principal-component analysis of the top 500 differentially expressed genes (DEGs). (c) Volcano plot showing differential expression between the two conditions; selected HOX- and glial-associated genes are labeled. (d) HOX gene heatmap showing the axial expression profile across the two conditions; asterisks indicate significantly different HOX genes identified by omnibus limma testing across groups with BH-adjusted *p* < 0.05. (e-f) Top enriched ARCHS4 tissue terms among genes elevated in diMNs 20DIM (e) or purMNs 20DIM (f), shown as horizontal bubble plots. (g-h) Top enriched GO terms among genes elevated in diMNs 20DIM (g) or purMNs 20DIM (h), grouped into Biological Process (BP), Cellular Component (CC), and Molecular Function (MF) categories. For panels e-h, enrichment was performed on genes elevated in each condition using the same differential-expression threshold; the x-axis indicates -log10(FDR), bubble size indicates overlapping gene number, and darker fill indicates stronger enrichment. n = 3 control iPSC lines per group.

Gene Ontology analysis further distinguished the two conditions, with diMNs enriched for developmental and proliferative programs (**Figure 2g**), whereas purMNs were enriched for synaptic signaling, postsynaptic membrane, ion transport, and glutamatergic receptor-associated terms (**Figure 2h**). Even at the matched 20DIM, the purified protocol yielded cultures that were more spinal and neuronally specialized than mixed diMN cultures.

Consistent with the enrichment analyses, examination of the genes contributing to these programs showed that purMNs expressed higher levels of caudal spinal and motor-neuron-associated genes, including *HOXA6, HOXA7, HOXA9, HOXC6, HOXC8, HOXC9, SLC6A5, GRM3, NTM, PCP4L1*, and *RELN*, together with neuronal maturation-associated genes such as *NEUROD2, NEUROD6, SLC30A3, RETREG1, LY6H, PCSK1N,* and *PNOC* (**Figure S1a-b**). purMNs also showed higher expression of genes contributing to postsynaptic, presynaptic/calcium, and cholinergic signaling programs, including *GRIA2, GRIA3, GRIA4, GRM3, SLC1A1, SLC1A2, GRM7, SYT7, CACNA1C, ADCY1, ADCY8, PRKCA, PRKCB, GNGT1, GNG7, KCNJ6, PLCB1,* and *CACNA1D* (**Figure S1c-e**). In contrast, diMNs retained higher expression of residual proliferation- and progenitor-associated markers, including *MKI67, TOP2A, CDK1, BIRC5, SOX1, NKX6-1,* and *ASCL1* (**Figure S1g-h**). Astroglial-lineage associated markers were more mixed, with *GFAP, PTGDS*, and *SLC22A17* being higher in purMNs, whereas *CXCL12* was higher in diMNs (**Figure S1f**), although this last gene is not only constrained to astrocyte lineage, and can also be a marker of stromal, endothelial, and epithelial cells^21^.

### Extended maturation further separates purified cultures and reinforces spinal axial identity

Having established that purMNs at 20DIM are already more caudalized and neuronally specialized than diMNs at 20DIM, we next tested whether additional time in maturation medium would further strengthen these spinal and synaptic features in purMNs. To do this, we extended only the purMN cultures to 32DIM and compared the transcriptomic outcomes with both diMNs and purMNs at 20DIM (**Figure 3a**). The protocols remained the same as before apart from the extended maturation period, which further separated purMNs 32DIM from purMNs 20DIM on principal-component analysis (**Figure 3b**).

**Figure 3.**
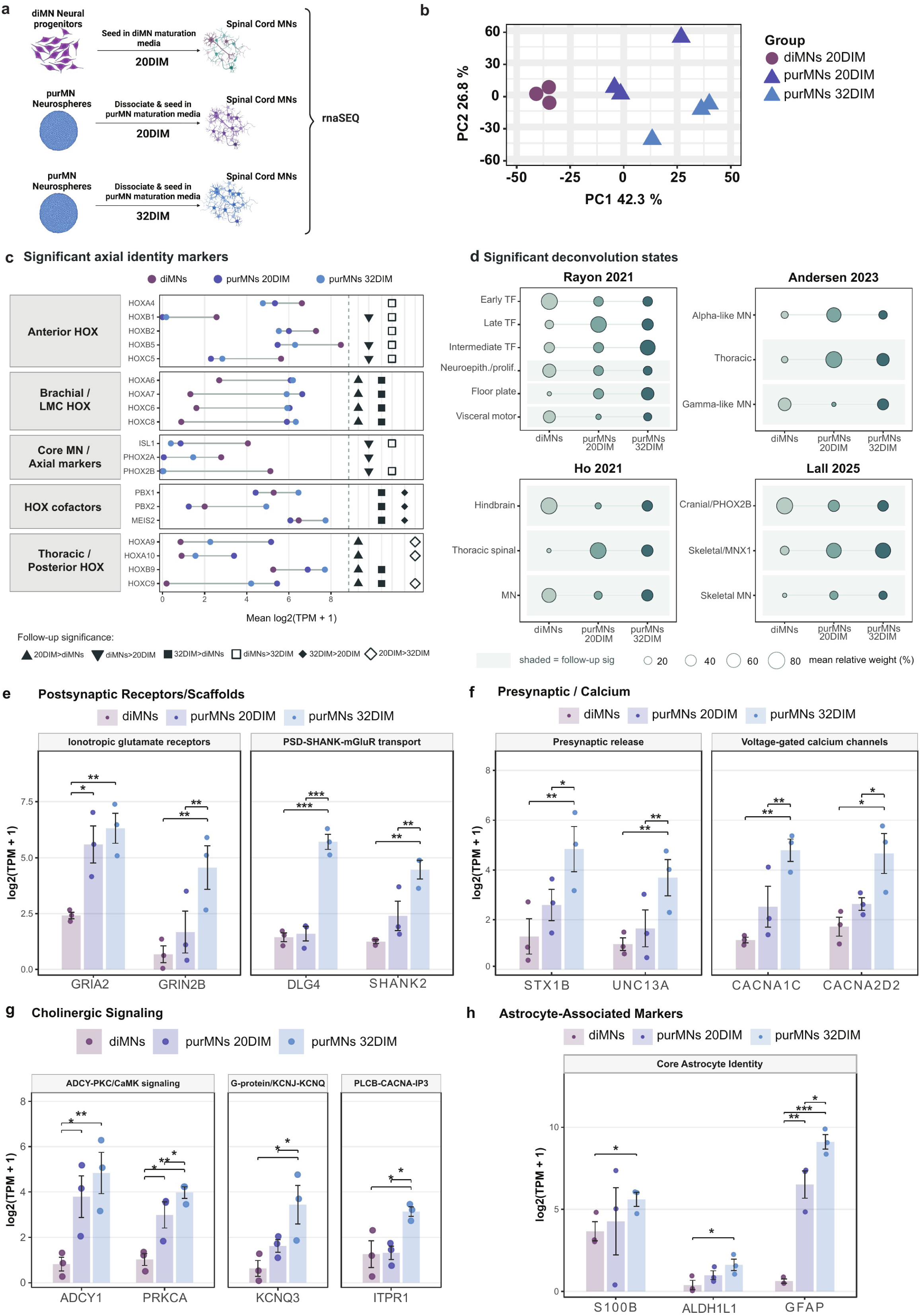
Extended maturation reinforces spinal identity and broadens synaptic, cholinergic, and astrocyte-associated transcriptional programs in purified cultures. (a) Experimental design for the three-group rnaSEQ comparison of diMNs 20DIM, purMNs 20DIM, and purMNs 32DIM. (b) Principal-component analysis showing separation by protocol and maturation duration. (c) Significant axial identity markers grouped into anterior HOX, brachial/LMC HOX, core motor neuron/axial markers, HOX cofactors, and thoracic/posterior HOX categories. Dots show mean log2(TPM + 1) expression for each group. Directional symbol columns at right indicate significant pairwise contrasts for purMNs 20DIM > diMNs, diMNs > purMNs 20DIM, purMNs 32DIM > diMNs, diMNs > purMNs 32DIM, purMNs 32DIM > purMNs 20DIM, and purMNs 20DIM > purMNs 32DIM. (d) Selected states from paper-guided reference-state scoring against (d1) Rayon et al., (d2) Andersen et al., (d3) Ho et al., and (d4) Lall et al. d1 and d2 provide *in vivo* developmental benchmarks, whereas d3 and d4 provide *in vitro* contextual references^17–20^. Bubble size and fill denote mean estimated relative weight, and row shading denotes states with BH-corrected follow-up pairwise contrast. (e) Postsynaptic receptor/scaffold genes. (f) Presynaptic/calcium genes. (g) Cholinergic signaling genes. (h) Astrocyte-associated genes. In panels c and e-h, n = 3 control iPSC lines per group and expression is shown as log2(TPM + 1). Bars show group means, error bars show SEM, and points show individual RNA-seq samples. Gene-level pairwise comparisons were performed using limma followed by BH correction; brackets and asterisks denote contrasts meeting BH-adjusted *p* < 0.05 and |logFC| >= 1. Significant HOX genes in panel c were identified by omnibus limma testing across groups with BH-adjusted *p* < 0.05, and directional symbols indicate significant pairwise limma contrasts meeting BH-adjusted *p* < 0.05 and |logFC| >= 1.

The three-group analysis reinforced the axial patterning trends seen in the two-group comparison. Significant axial identity markers showed that diMNs at 20DIM retained a more anterior/cervical profile, whereas purMNs at 20DIM and purMNs at 32DIM aligned more strongly with broader spinal programs featuring brachial/thoracic associated HOX expression and posterior axial markers (**Figure 3c**).

To place these transcriptomic differences in context, we compared the bulk RNA-seq profiles with four published single-cell reference frameworks, separating *in vivo* developmental benchmarks of the developing human spinal cord^17,18^, from *in vitro* iPSC-derived motor neurons^19,20^ (**Figure 3d**; **Figure S2**). Although each framework used distinct state definitions, the overall direction of change was consistent, with diMNs at 20DIM aligned more strongly with anterior, cranial/ *PHOX2B*, hindbrain-like, or progenitor-associated states, while purMNs at 20DIM and purMNs at 32DIM aligned more strongly with spinal, thoracic, skeletal motor neuron, and postmitotic neuron-associated programs.

### Pairwise enrichment and gene-family analyses identify transcriptional programs associated with differentiation protocol and purMN maturation

To characterize the biological programs distinguishing diMNs at 20DIM, purMNs at 20DIM, and purMNs at 32DIM, tissue, KEGG, and Gene Ontology (GO) enrichment analyses were performed separately for genes increased in each directional pairwise comparison (**Figure S4**). Compared with diMNs, purMNs at 32DIM were enriched for central nervous system and motor neuron-associated tissue terms, together with synaptic and axon-related pathways including glutamatergic synapse, calcium signaling, cAMP signaling, and axon guidance (**Figure S4a-b**). GO analysis similarly identified enrichment for synapse organization, trans-synaptic signaling, postsynaptic specialization, ion-channel activity, and axonogenesis (**Figure S4c**). Compared with purMNs at 20DIM, purMNs at 32DIM were enriched for small GTPase signaling, cell-junction assembly, neuronal projection development, axonogenesis, and synapse organization (**Figure S4d**). Conversely, genes more highly expressed in diMNs or purMNs at 20DIM than in purMNs at 32DIM were enriched for mitochondrial respiration, oxidative phosphorylation, mitochondrial translation, and respiratory-chain complexes, with DNA-replication terms additionally enriched in diMNs (**Figure S4e-f**).

We next examined representative transcripts grouped into postsynaptic, presynaptic and signaling families (**Figure 3e**), with expanded details shown in **Figure S3a-c**. The most prominent changes were observed in purMNs at 32DIM, which compared to 20DIM showed even higher expression of multiple ionotropic glutamate receptor subunits, *PSD/SHANK*-associated genes, presynaptic release genes, and voltage-gated calcium-channel components (**Figure 3e-f; Figure S3a-b**). Cholinergic pathway-related signaling genes, including adenylate cyclases, *PKC/CaMK*-related components, G-protein/*KCNJ-KCNQ* modules, and *PLCB-CACNA-IP3* signaling genes, were also elevated (**Figure 3g; Figure S3c**).

In parallel, prolonged maturation was associated with increased expression of astrocyte-associated markers (*GFAP, S100B, ALDH1L1, and SLC1A2*), developmental regulators, such as *SOX9*, reactive/signaling-associated genes, like *CXCL12*, and secreted or membrane-associated genes (**Figure 3h; Figure S3d**). Together, gene enrichments indicate that prolonged maturation remodels both synaptic and astroglial-associated transcriptional programs rather than simply amplifying a generic neuronal signal.

### High-purity protocol yields cultures enriched in proteins for motor neurons and with a distinct glial background

We next assessed how time in culture affected MN numbers based on immunofluorescence and quantification of canonical MN markers. The diMN protocol at 20 DIM yielded a substantially smaller fraction of cells that were ChAT^+^ (27.94%*±*7.21), MNX1^+^ (45.23%*±*10.38), and ISL1/2^+^ (18.23%*±*2.78) (**Figure 4a-b,g**) in contrast to the purMN protocol at 32 DIM that yielded cultures strongly enriched for ChAT^+^ (81.54%*±*5.02), MNX1^+^ (81.27%*±*3.62), and ISL1/2^+^ (76.17%*±*8.14) cells (**Figure 4d-e,g**). The protocols also yielded a distinct profile of glial markers. diMN cultures showed S100p (radial immunoreactivity (49.44%*±*10.42) in flat cells with large nuclei but no detectable glial fibrillary acidic protein (GFAP), a canonical astrocyte marker (**Figure 4c,g**), suggesting the presence of a non-neuronal glial/progenitor-associated population rather than mature GFAP-positive astrocytes. In contrast, purMNs lacked this broad S100p-positive component but contained a small GFAP-positive (7.32%*±*2.79) population with distinct morphology (**Figure 4f,g**).

**Figure 4.**
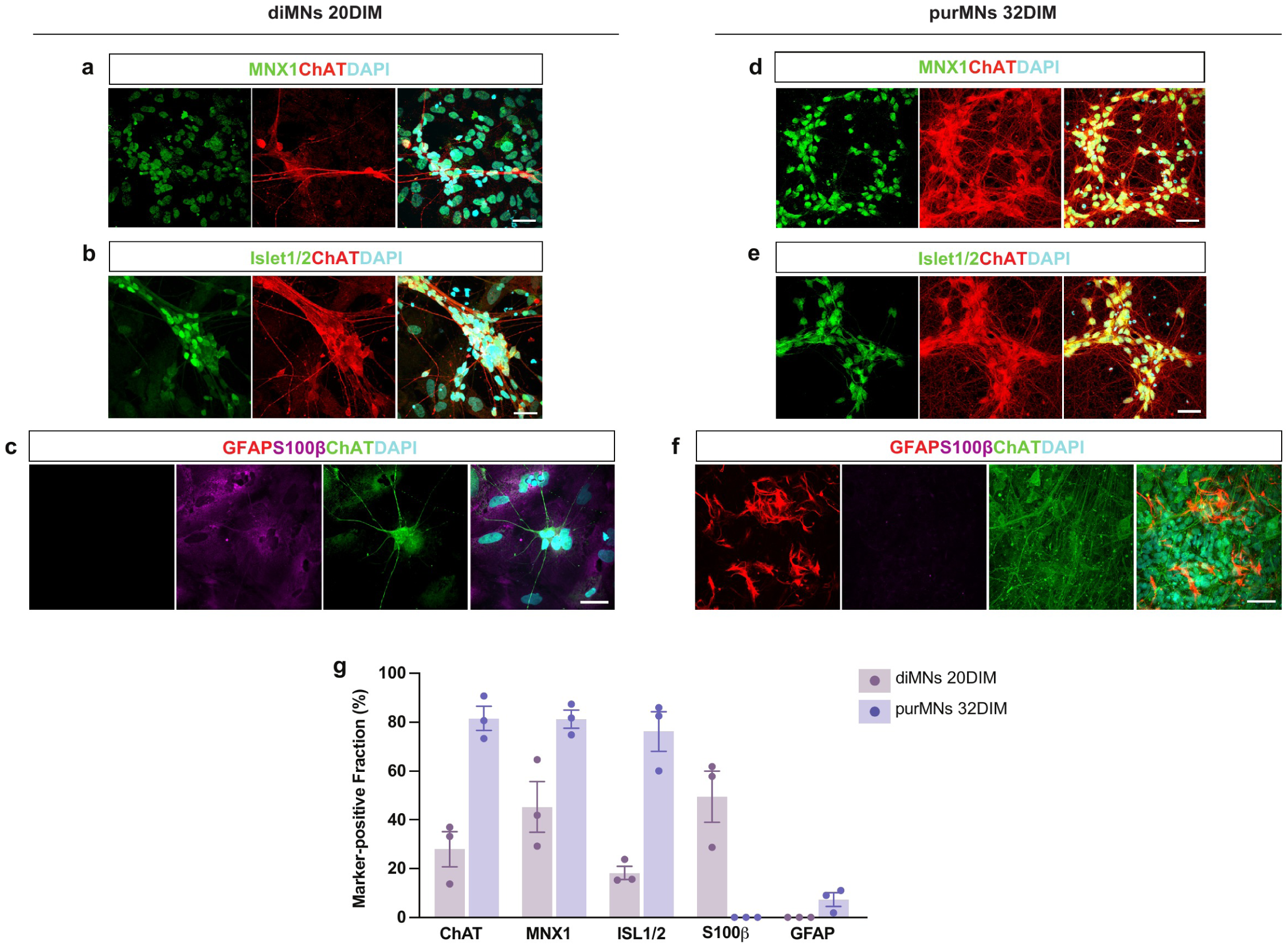
High-purity differentiation yields motor neuron-enriched cultures with distinct glial-associated backgrounds. (a-c) Representative diMN immunofluorescence images at 20DIM. (d-f) Representative purMN immunofluorescence images at 32DIM. (a, d) MNX1 (green), ChAT (red), and DAPI (cyan). (b, e) ISL1/2 (green), ChAT (red), and DAPI (cyan). (c, f) GFAP (red), S100B (magenta), ChAT (green), and DAPI (cyan). Individual channels and merged images are shown; DAPI was pseudocolored cyan. (g) Quantification of ChAT-, MNX1-, and ISL1/2-positive nuclei as percentages of total DAPI-positive nuclei, and S100B-and GFAP-positive areas as percentages of total image area. (g) Quantification for MN and astroglial-related markers summarizing the percentage of marker positive cells. Quantification is based on n=3 control iPSC lines, with 3 independent differentiations per line per protocol. Points represent the average expression for each line, bars show mean, and error bars show SEM. Multiple non-overlapping fields were analyzed per coverslip and averaged within each biological replicate. Scale bars, 30 μm.

### Purified motor neurons form denser neurite networks and show stronger structural neuromuscular engagement in microfluidic co-culture

Given that spinal MNs connect to muscle *in vivo* as a neuromuscular junction (NMJ), we next asked whether the transcriptomic maturation associated with the purified protocol was accompanied by structural differences in neuron-muscle co-culture. To address this question, we utilized a microfluidic NMJ-chip with 3 chambers, into which we seeded either iPSC-derived purMNs or diMNs into 2 channels, followed at 14 days with seeding a central compartment with iPSC-derived skeletal muscle progenitor cells (SMPCs) (**Figure 5a**). In all cases, the same control iPSC line was used. The neurites extending from diMNs and purMNs were stained for neurofilament heavy and these were overlayed with skeletonized/thickness using the Fiji program, which showed that the purMNs formed visibly denser neurite networks than diMNs (**Figure 5b-c**). Quantitative analysis showed that purMN cultures had greater branch length per area, more branch junctions, and more endpoints (**Figure 5d-f**), whereas mean branch length was not detectably changed (**Figure 5g**). At the same time, purMNs showed reduced weighted median thickness and reduced 90th-percentile neurite thickness (**Figure 5h-i**), indicating that the denser network consisted of finer, more highly branched projections rather than simply thicker processes. To assess the connection between MNs and SMPCs in the central compartment, motor neuron axons were stained for neurofilament heavy, while SMPCs were stained for myosin heavy chain and acetylcholine receptor (AChR) clusters were labeled with alpha-bungarotoxin. Axons from both MN protocols contacted alpha-bungarotoxin-labeled AChR clusters, but the purMN signal appeared greater (**Figure 5j-k**). Quantification showed that purMNs generated significantly more AChR clusters and more innervated clusters compared to diMNs (**Figure 5l-m**). These phenotypes are consistent with the synaptic and axon-remodeling programs observed in purMN cultures.

**Figure 5.**
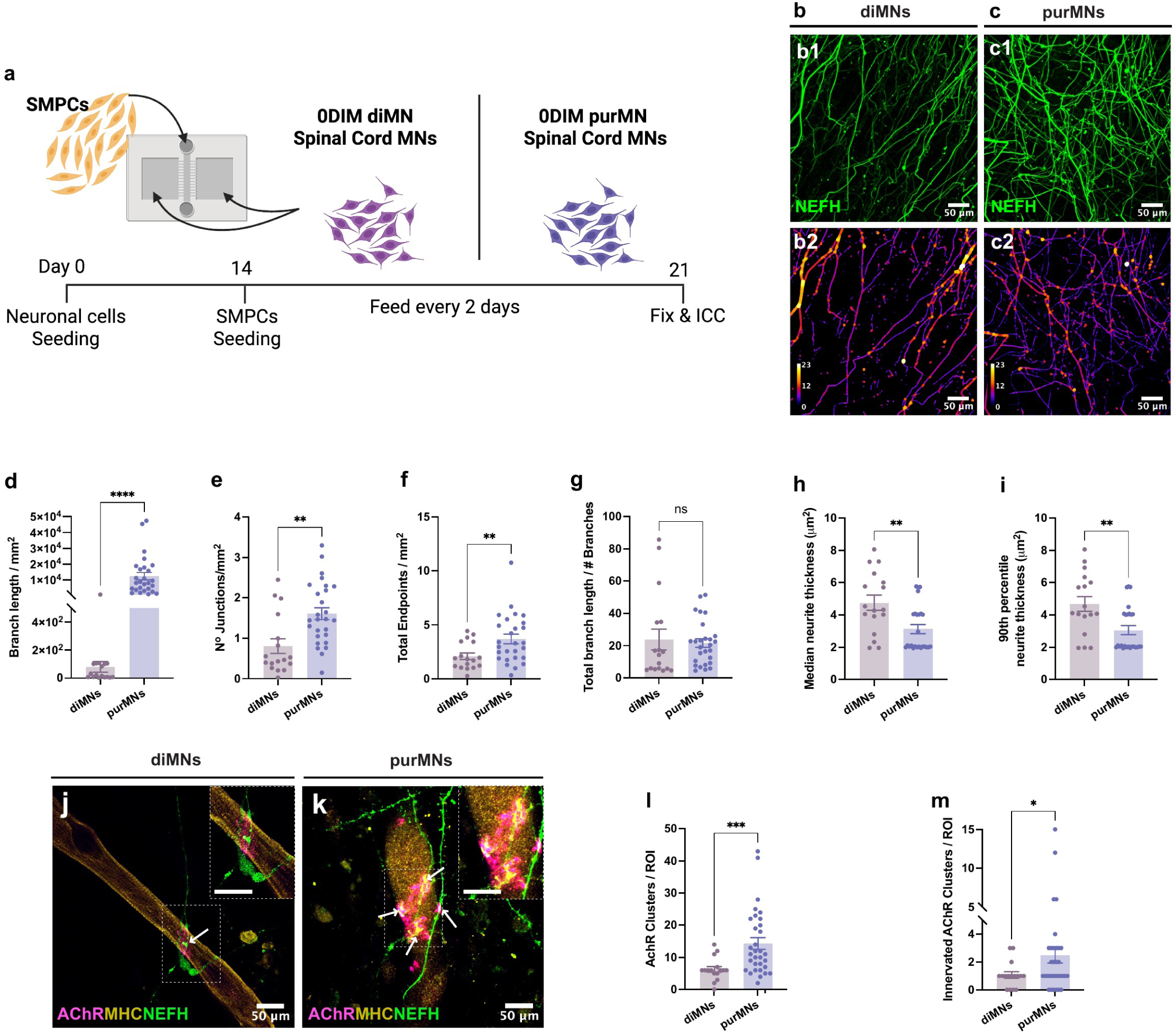
Purified motor neurons form denser neurite networks and show greater structural neuromuscular engagement in microfluidic co-culture. (a) Schematic of the Xona microfluidic NMJ-chip workflow using one control iPSC line. diMNs or purMNs were seeded into the lateral neuronal compartments. SMPCs derived from one iPSC line was seeded into the central muscle compartment after neurite extension through the microgrooves. (b-c) Representative Neurofilament Heavy (NEFH)-stained distal neurite images and corresponding skeletonized local-thickness overlays for diMN (b2) and purMN (c2) co-cultures. Local-thickness overlays are displayed using the same color scale across conditions; color denotes thickness in pixels. Scale bars, 50 *μ*m. (d-i) Quantification of neurite morphology, including total branch length per mm^2^ (d), branch junctions per mm^2^ (e), endpoints per mm^2^ (f), mean branch length (g), weighted median neurite thickness (h), and 90th-percentile neurite thickness (i). (j-k) Representative images showing NEFH-positive motor axons, myosin heavy chain-positive myotubes, and alpha-bungarotoxin-labeled AChR clusters. Scale bars, 50 *μ*m. (l-m) Quantification of total AChR clusters per ROI (l) and innervated AChR clusters per ROI (m), defined as AChR clusters located within a muscle fiber and in contact with a neurite. For all quantified panels, points represent ROIs from n = 3 independent chips per condition, bars show mean, and error bars show SEM. Statistical significance was determined by unpaired two-tailed t-tests with Welch’s correction. \**p* < 0.05, \*\**p* < 0.01, \*\*\**p* < 0.001; ns, not significant.

## Discussion

This study identifies protocol choice and time in maturation medium as major determinants of transcriptomic state in human iPSC-derived spinal motor neuron cultures. Although both differentiation strategies follow the broad developmental logic of neural induction, caudalization, and ventralization, prior work shows that motor neuron output is highly sensitive to the timing and combination of patterning cues, including RA, SHH-pathway activation, WNT activation, and BMP/TGF-beta inhibition^12,22^. While we compared the two workflows at the same days in maturation media, it’s important to note that the protocols differ in total differentiation age and diverge at several protocol levels, including the timing and composition of early neural induction, the use of DMH1-based BMP inhibition during neuroepithelial induction in the purMN workflow, additional neuroepithelial, motor neuron progenitor, and neurosphere stages in the purMN workflow, and trophic factor delivery^2,23–26^. The upstream differentiation and patterning history, culture format, and overall protocol duration likely act together to influence motor neuron yield, axial identity, maturation state, and residual non-neuronal programs, rather than reflecting the effect of simply the protocol days in maturation. Given these overall protocol differences, it is important knowledge for the field to compare the published protocols and clarify their motor neuron output. The purMN workflow yielded more purified cultures with stronger MNX1, ISL1/2, and ChAT motor neuron signatures together with a small GFAP-positive compartment, whereas the diMN workflow showed mixed cultures that retained a broader S100p-positive flat-cell background lacking GFAP in addition to broader progenitor-like and non-motor-neuron-associated features. This distinction is consistent with the original high-purity spinal motor neuron progenitor strategy described by Du et al. (2015)^2^, which was designed to generate lineage-restricted motor neuron progenitors efficiently, and contrasts with large-scale mixed differentiation approaches that were optimized in part for scalability and reproducibility across many patient lines^3,6^.

At matched 20DIM, purMN cultures were already clearly distinct from diMNs in composition, axial identity, and neuronal specialization. The purified cultures showed stronger caudal HOX expression and broader enrichment of neuronal and synaptic programs, whereas diMNs retained stronger developmental and progenitor-associated signatures. These findings support the idea that differences between protocols are not simply quantitative differences in motor neuron yield, but also qualitative differences in axial specification and lineage resolution. This matters because rostrocaudal patterning is tightly linked to spinal motor neuron subtype identity and function^17^. Our findings are also consistent with prior transcriptomic work showing that iPSC-derived spinal motor neuron cultures are heterogeneous and contain biologically meaningful subtype variation that cannot be captured by a small number of canonical markers alone^27^.

A central finding of this study is that extended maturation of the purified cultures is not a passive holding period but instead promotes additional transcriptional remodeling toward a more mature motor neuron state. Because purMNs at 20DIM already displayed a more caudalized and neuronally specialized profile than diMNs, we specifically asked whether additional time in maturation medium would further reinforce these spinal and synaptic features in the purified condition. The extension of purMNs from 20DIM to 32DIM was associated with stronger expression of glutamatergic receptor subunits, postsynaptic scaffold genes, presynaptic release machinery, calcium-signaling components, and cholinergic pathway-related signaling genes, together with broader enrichment of synapse organization, membrane excitability, and trans-synaptic communication terms. This profile aligns with developmental studies showing that motor neuron maturation involves progressive and stage-specific transcriptional transitions rather than simple upregulation of a few terminal differentiation markers^28^. It is also consistent with reports that long-term culture conditions are required to stabilize more mature spinal motor neuron features *in vitro*^29,30^.

The paper-guided deconvolution analyses provides an independent line of support for these conclusions. Across four distinct single-cell reference frameworks, diMNs aligned more strongly with anterior, cranial, hindbrain-like, gamma-like, or progenitor-associated states, whereas purMN cultures aligned more strongly with spinal, thoracic, skeletal motor neuron, and post-mitotic neuron-associated programs. The fact that this general direction was reproduced across multiple references is important because each atlas emphasizes different biological dimensions, including rostrocaudal identity, dorsoventral patterning, temporal neuronal code, glial lineage emergence, and motor neuron subtype diversification^17–20^. Together, these comparisons argue that the purMN workflow does not simply increase motor neuron identity in a generic sense but instead shifts cultures toward a more spinally patterned and post-mitotic transcriptional state. At the same time, these outputs should be interpreted as relative alignment to external reference programs rather than direct estimates of cell fraction. Another important point is that prolonged maturation was accompanied not only by stronger synaptic and signaling-associated programs, but also by the emergence of some astroglial-associated transcriptional features. These results are consistent with the proteomic characterization of cells generated by the purMN protocol^31^. We do not interpret this as a loss of motor neuron identity. Rather, these findings suggest that the 32DIM cultures retain strong motor neuron features while also showing a modest increase in support-cell or lineage-adjacent transcriptional signatures. This interpretation is consistent with prior work showing that extended iPSC neuronal culture promotes morphological, electrophysiological, and transcriptional maturation, including synaptic pathway^32,33^. It is also consistent with single-cell and large-scale iPSC-derived motor neuron studies showing that motor neuron-directed differentiations can retain minority interneuron, progenitor, glial/progenitor-like, or S100p-positive non-neuronal populations that contribute to bulk transcriptomic variation^3,34^. Therefore, the 32DIM state should not be viewed simply as a purer version of the 20DIM state; instead, longer maturation may improve access to synaptic programs while also increasing the low presence of of non-motor neuron -associated signals.

The structural microfluidic co-culture data showed that purMN cultures formed denser and more highly branched distal axonal networks and showed stronger AChR-cluster engagement than diMN cultures, indicating that the identified molecular differences are accompanied by measurable morphological consequences in neuron-muscle assays. This is in line with the view that more mature and more faithfully patterned spinal motor neurons should also show improved distal projection behavior and target engagement. The effects of enhanced maturation are also relevant in the context of a recent organ-chip model of ALS, which developed disease phenotypes when patient iPSC-derived spinal MNs were cultured in conditions that promoted further differentiation and maturation^20^.

While this study shows that protocol choice and maturation can lead to clear and important differences in motor neuron cultures, there do exist some limitations. Given that the transcriptomic analyses were performed in bulk, lineage complexity and cell state heterogeneity were inferred indirectly rather than measured at the single-cell level. Additionally, only three iPSC lines were used, and the NMJ neurite analyses were performed with a single line. Therefore, while all experiments were performed across multiple differentiation studies, future studies should include additional lines. Finally, the deconvolution analyses depend on the structure and resolution of the external reference datasets and therefore should be interpreted conservatively. Even with these limitations, the benchmarking framework presented here provides a practical basis for selecting differentiation conditions according to the intended biological questions and experimental needs. We have demonstrated that the purMN protocol provides a stronger and more interpretable motor neuron state than the mixed diMN protocol. The shorter diMN protocol does generate motor neurons, but at a lower number and maturity. However, this protocol can still benefit large scale consortium studies. In contrast, the extended purMN workflow with increased maturation may be best for studies wanting to interrogate spinal identity, synaptic maturation, cholinergic signaling, distal axonal architecture, or neuromuscular engagement. These findings can help guide the field when designing experiments to study motor neuron development, health and disease.

## Acknowledgements

This work was supported by the California Institute for Regenerative Medicine (CIRM) Scholar Training Program EDUC4-12751 (A.S., V.Z., M.F.). We thank members of the Svendsen laboratory for their assistance and Dr. Soshana Svendsen for her review of the manuscript. We thank the members of the Cedars-Sinai Applied Genomics, Computation & Translational Core for their assistance in RNA sequencing.

## Author Contributions

Conceptualization: A.S., C.N.S. Methodology: A.S., V.Z., J.V, M.F., S.B. Investigation: A.S., V.Z., M.F. Formal analysis: A.S., V.Z., S.B. Data curation: A.S. Visualization: A.S. Supervision: C.N.S. Funding acquisition: C.N.S., A.S., Writing—original draft: A.S. Writing—review and editing: A.S., V.Z., C.N.S.

## Lead Contact

Further information and requests for resources should be directed to and will be fulfilled by the Lead Contact, Clive N. Svendsen.

## Materials Availability

This study did not generate new unique reagents. Human iPSC lines are available from the Cedars-Sinai Biomanufacturing Center subject to standard material transfer agreements.

## Data and Code Availability

The bulk RNA-seq data generated in this study have been deposited in NCBI Gene Expression Omnibus and will be available under accession number GEO: GSE346255. Processed gene-level count and TPM matrices are included with the GEO submission.

## Declaration of generative AI and AI-assisted technologies in the manuscript preparation process

During the preparation of this work, the author(s) used Codex (OpenAI) to assist with organizing and revising the manuscript text and figure legends, and the development and documentation of bioinformatic analysis code. The author(s) reviewed and edited the output as needed and take full responsibility for the content of the published article. Codex was not used to generate primary experimental data.

