## Supplementary Figures and Tables for "Differentiation protocol and maturation shape the axial identity and synaptic state in human iPSC-derived spinal motor neurons"

#### Figure S1

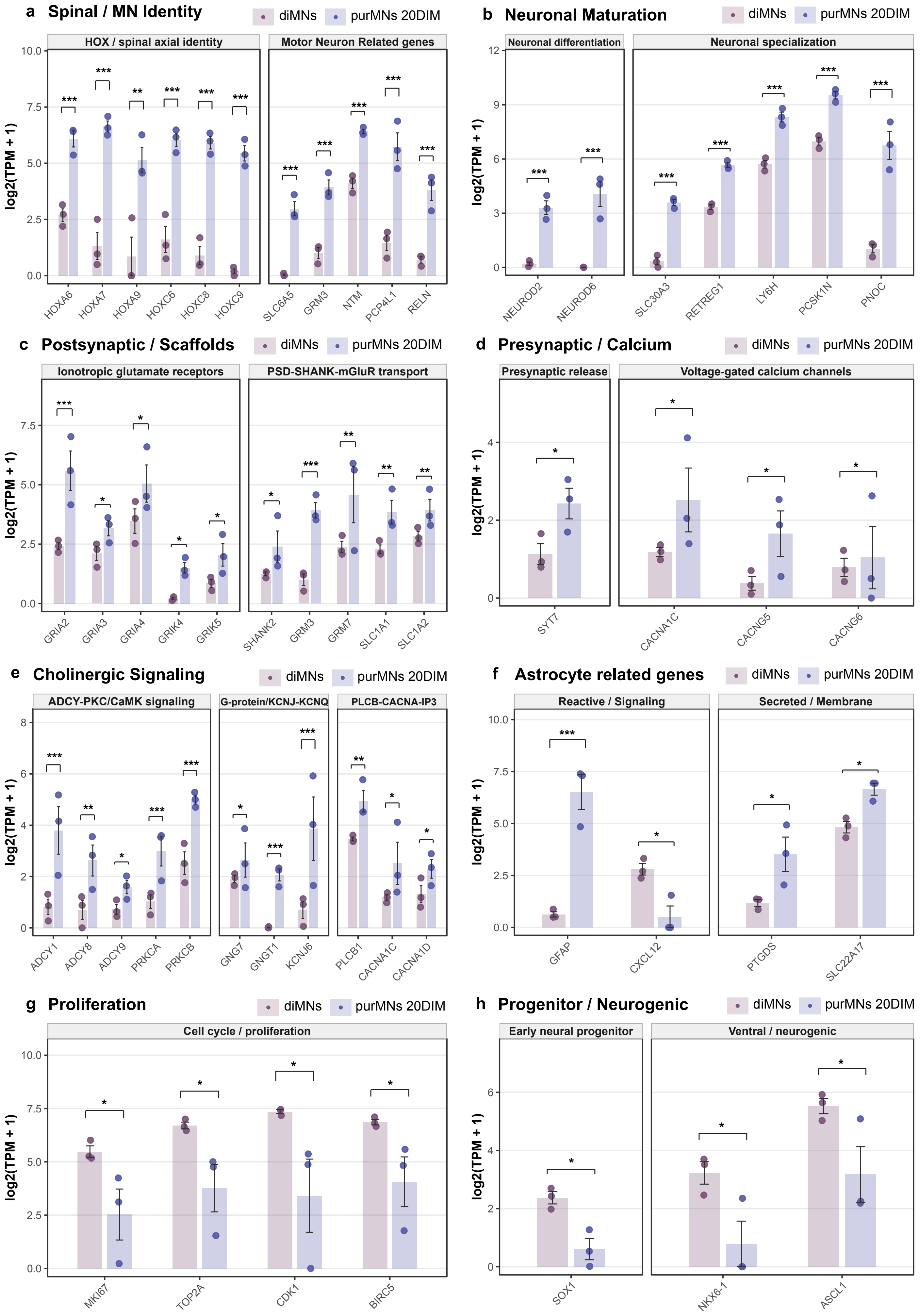

a Spinal / MN Identity
diMNs
purMNs 20DIM
HOX / spinal axial identity
Motor Neuron Related genes
log2(TPM + 1)
HOXA6
HOXA7
HOXA9
HOXC6
HOXC8
HOXC9
SLC6A5
GRM3
NTM
PCP4L1
RELN
b Neuronal Maturation
Neuronal differentiation
Neuronal specialization
NEUROD2
NEUROD6
SLC30A3
RETREG1
LY6H
PCSK1N
PNOC
c Postsynaptic / Scaffolds
Ionotropic glutamate receptors
PSD-SHANK-mGluR transport
GRIA2
GRIA3
GRIA4
GRIK4
GRIK5
SHANK2
GRM3
GRM7
SLC1A1
SLC1A2
d Presynaptic / Calcium
Presynaptic release
Voltage-gated calcium channels
SYT7
CACNA1C
CACNG5
CACNG6
e Cholinergic Signaling
ADCY-PKC/CaMK signaling
G-protein/KCNJ-KCNQ
PLCB-CACNA-IP3
ADCY1
ADCY8
ADCY9
PRKCA
PRKCB
GNG7
GNGT1
KCNJ6
PLCB1
CACNA1C
CACNA1D
f Astrocyte related genes
Reactive / Signaling
Secreted / Membrane
GFAP
CXCL12
PTGDS
SLC22A17
g Proliferation
Cell cycle / proliferation
MKI67
TOP2A
CDK1
BIRC5
h Progenitor / Neurogenic
Early neural progenitor
Ventral / neurogenic
SOX1
NKX6-1
ASCL1

**Figure S1. Targeted two-group marker panels highlighting the genes underlying the major programs differentiating diMNs and purMNs at matched 20DIM, related to Figure 2.**

(a) Spinal / motor neuron identity markers. (b) Neuronal maturation-associated markers. (c) Postsynaptic receptor and scaffold markers. (d) Presynaptic release and voltage-gated calcium channel markers. (e) Cholinergic pathway signaling markers. (f) Astrocyte-associated markers. (g) Proliferation markers. (h) Progenitor / neurogenic markers. All panels show expression in the comparing diMNs 20DIM versus purMNs 20DIM as log2(TPM + 1). Bars indicate group means, error bars indicate SEM, and points indicate individual samples. Genes were selected to represent the major biological programs identified in the analysis. Brackets indicate pairwise significance from the dedicated two-group limma analysis using a relaxed BH-adjusted threshold (* adjusted $p < 0.25$, ** adjusted $p < 0.10$, *** adjusted $p < 0.05$).

### Figure S2

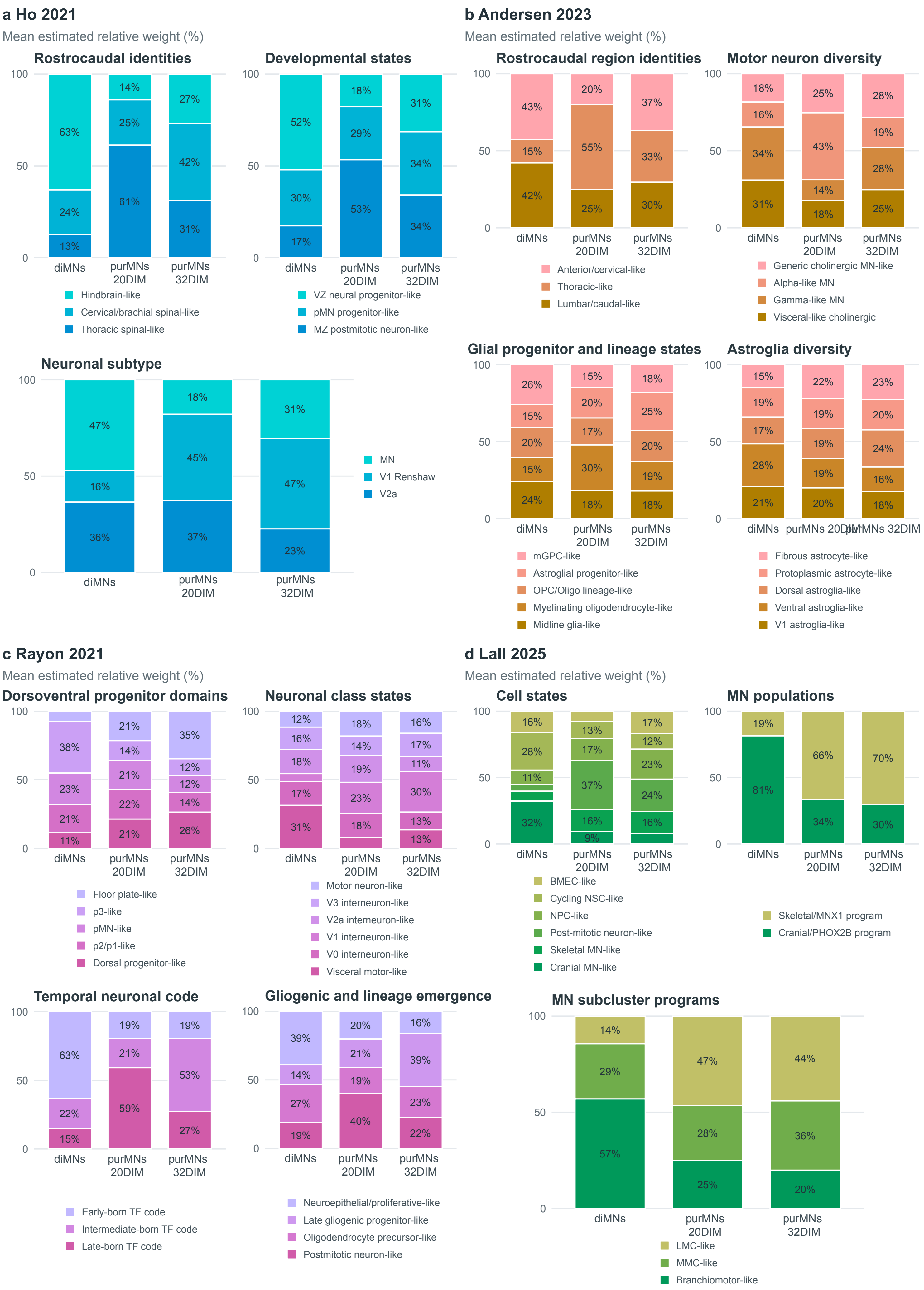
a Ho 2021
Mean estimated relative weight (%)
Rostrocaudal identities
100
50
0
63%
24%
13%
14%
25%
61%
27%
42%
31%
diMNs
purMNs 20DIM
purMNs 32DIM
Hindbrain-like
Cervical/brachial spinal-like
Thoracic spinal-like
Developmental states
52%
30%
17%
18%
29%
53%
31%
34%
34%
VZ neural progenitor-like
pMN progenitor-like
MZ postmitotic neuron-like
Neuronal subtype
47%
16%
36%
18%
45%
37%
31%
47%
23%
MN
V1 Renshaw
V2a
b Andersen 2023
Mean estimated relative weight (%)
Rostrocaudal region identities
43%
15%
42%
20%
55%
25%
37%
33%
30%
Anterior/cervical-like
Thoracic-like
Lumbar/caudal-like
Motor neuron diversity
18%
16%
34%
31%
25%
43%
14%
18%
28%
19%
28%
25%
Generic cholinergic MN-like
Alpha-like MN
Gamma-like MN
Visceral-like cholinergic
Glial progenitor and lineage states
26%
15%
20%
15%
24%
15%
20%
17%
30%
18%
18%
25%
20%
19%
18%
mGPC-like
Astroglial progenitor-like
OPC/Oligo lineage-like
Myelinating oligodendrocyte-like
Midline glia-like
Astroglia diversity
15%
19%
17%
28%
21%
22%
19%
19%
19%
20%
23%
20%
24%
16%
18%
Fibrous astrocyte-like
Protoplasmic astrocyte-like
Dorsal astroglia-like
Ventral astroglia-like
V1 astroglia-like
c Rayon 2021
Mean estimated relative weight (%)
Dorsoventral progenitor domains
38%
23%
21%
11%
21%
14%
21%
22%
21%
35%
12%
12%
14%
26%
Floor plate-like
p3-like
pMN-like
p2/p1-like
Dorsal progenitor-like
Neuronal class states
12%
16%
18%
17%
31%
18%
14%
19%
23%
18%
16%
17%
11%
30%
13%
13%
Motor neuron-like
V3 interneuron-like
V2a interneuron-like
V1 interneuron-like
V0 interneuron-like
Visceral motor-like
Temporal neuronal code
63%
22%
15%
19%
21%
59%
19%
53%
27%
Early-born TF code
Intermediate-born TF code
Late-born TF code
Gliogenic and lineage emergence
39%
14%
27%
19%
20%
21%
19%
40%
16%
39%
23%
22%
Neuroepithelial/proliferative-like
Late gliogenic progenitor-like
Oligodendrocyte precursor-like
Postmitotic neuron-like
d Lall 2025
Mean estimated relative weight (%)
Cell states
16%
28%
11%
32%
13%
17%
37%
16%
9%
17%
12%
23%
24%
16%
BMEC-like
Cycling NSC-like
NPC-like
Post-mitotic neuron-like
Skeletal MN-like
Cranial MN-like
MN populations
19%
81%
66%
34%
70%
30%
Skeletal/MNX1 program
Cranial/PHOX2B program
MN subcluster programs
14%
29%
57%
47%
28%
25%
44%
36%
20%
LMC-like
MMC-like
Branchiomotor-like

**Figure S2. Four independent paper-guided deconvolution frameworks converge on a spinal/postmitotic shift in purified cultures, related to Figure 3.**

Each quadrant shows one paper-guided deconvolution framework, with mean estimated relative weights displayed as stacked bars and percentages shown within segments. (a) Rayon 2021 resolves dorsoventral progenitor domains, neuronal class states, temporal neuronal codes, and gliogenic/lineage-emergence states. (b) Andersen 2023 resolves rostrocaudal regional identities, motor neuron diversity, glial progenitor/lineage states, and astroglial diversity. (c) Ho 2021 resolves rostrocaudal identity, developmental states, and neuronal subtype states. (d) Lall 2025 resolves cell states, motor neuron population programs, and motor neuron subcluster programs. Across frameworks, diMNs align more strongly with anterior, cranial, hindbrain-like, or progenitor-associated programs, whereas purMNs 20DIM and purMNs 32DIM align more strongly with spinal, thoracic, skeletal motor neuron, and postmitotic neuron-associated programs. For each deconvolution framework, state weights were compared across diMNs 20DIM, purMNs 20DIM, and purMNs 32DIM using n = 3 control iPSC lines per group. Omnibus one-way ANOVA was followed by BH correction across states. Values are shown as mean relative state weights and should be interpreted as framework-aligned relative weights rather than direct cell-fraction estimates.

#### Figure S3

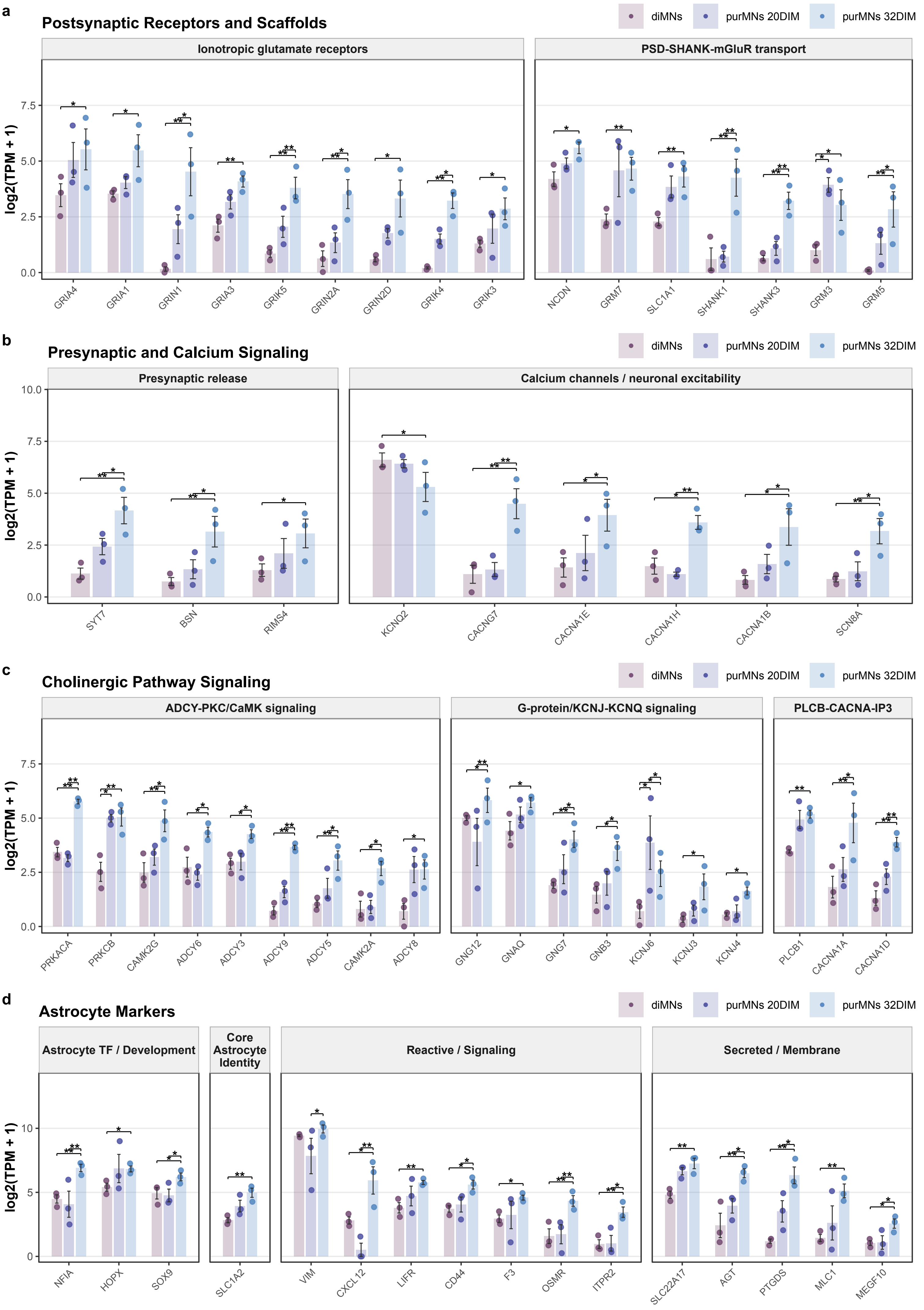

a
Postsynaptic Receptors and Scaffolds
diMNs
purMNs 20DIM
purMNs 32DIM
Ionotropic glutamate receptors
PSD-SHANK-mGluR transport
log2(TPM + 1)
GRIA4
GRIA1
GRIN1
GRIA3
GRIK5
GRIN2A
GRIN2D
GRIK4
GRIK3
NCDN
GRM7
SLC1A1
SHANK1
SHANK3
GRM3
GRM5
b
Presynaptic and Calcium Signaling
Presynaptic release
Calcium channels / neuronal excitability
SYT7
BSN
RIMS4
KCNQ2
CACNG7
CACNA1E
CACNA1H
CACNA1B
SCN8A
c
Cholinergic Pathway Signaling
ADCY-PKC/CaMK signaling
G-protein/KCNJ-KCNQ signaling
PLCB-CACNA-IP3
PRKACA
PRKCB
CAMK2G
ADCY6
ADCY3
ADCY9
ADCY5
CAMK2A
ADCY8
GNG12
GNAQ
GNG7
GNB3
KCNJ6
KCNJ3
KCNJ4
PLCB1
CACNA1A
CACNA1D
d
Astrocyte Markers
Astrocyte TF / Development
Core Astrocyte Identity
Reactive / Signaling
Secreted / Membrane
NFIA
HOPX
SOX9
SLC1A2
VIM
CXCL12
LIFR
CD44
F3
OSMR
ITPR2
SLC22A17
AGT
PTGDS
MLC1
MEGF10

**Figure S3. Expanded synaptic, cholinergic, and astrocyte-associated gene modules underlying purified motor neuron maturation, related to Figure 3.**

(a) Postsynaptic receptor and scaffold genes, grouped into ionotropic glutamate receptors and *PSD-SHANK-mGluR* transport modules. (b) Presynaptic release genes and voltage-gated calcium-channel genes. (c) Cholinergic pathway signaling genes grouped into *ADCY-PKC/CaMK*, *G-protein/KCNJ-KCNQ*, and *PLCB-CACNA-IP3* modules. (d) Astrocyte-associated genes grouped into astrocyte Transcription factor (TF)/development, core astrocyte identity, reactive/signaling-associated, and secreted/membrane modules. Bars show group means for diMNs 20DIM, purMNs 20DIM, and purMNs 32DIM; error bars show SEM; points show individual RNA-seq samples; n = 3 control iPSC lines per group. Pairwise comparisons were performed using limma on log2(TPM + 1) values followed by BH correction. Asterisks indicate contrasts meeting both the indicated adjusted-p-value threshold and |logFC| >= 1: * adjusted $p < 0.05$, ** adjusted $p < 0.01$, *** adjusted $p < 0.001$.

#### Figure S4

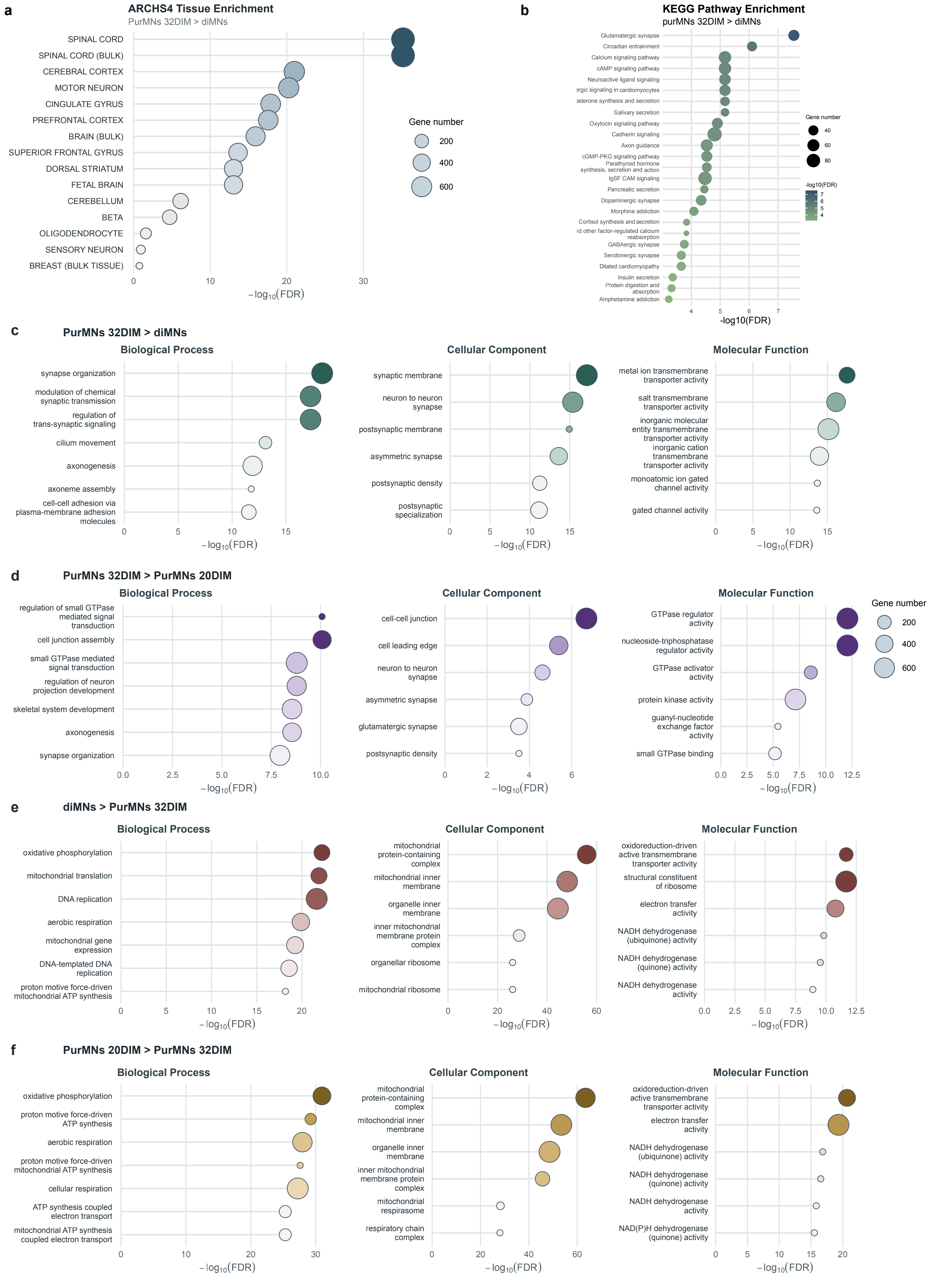
a
ARCHS4 Tissue Enrichment
PurMNs 32DIM > diMNs
SPINAL CORD
SPINAL CORD (BULK)
CEREBRAL CORTEX
MOTOR NEURON
CINGULATE GYRUS
PREFRONTAL CORTEX
BRAIN (BULK)
SUPERIOR FRONTAL GYRUS
DORSAL STRIATUM
FETAL BRAIN
CEREBELLUM
BETA
OLIGODENDROCYTE
SENSORY NEURON
BREAST (BULK TISSUE)
0
10
20
30
$-\log_{10}(FDR)$
Gene number
200
400
600
b
KEGG Pathway Enrichment
purMNs 32DIM > diMNs
Glutamatergic synapse
Circadian entrainment
Calcium signaling pathway
cAMP signaling pathway
Neuroactive ligand signaling
ergic signaling in cardiomyocytes
osterone synthesis and secretion
Salivary secretion
Oxytocin signaling pathway
Cadherin signaling
Axon guidance
cGMP-PKG signaling pathway
Parathyroid hormone synthesis, secretion and action
IgSF CAM signaling
Pancreatic secretion
Dopaminergic synapse
Morphine addiction
Cortisol synthesis and secretion
nd other factor-regulated calcium reabsorption
GABAergic synapse
Serotonergic synapse
Dilated cardiomyopathy
Insulin secretion
Protein digestion and absorption
Amphetamine addiction
4
5
6
7
-log10(FDR)
Gene number
40
60
80
-log10(FDR)
7
6
5
4
c
PurMNs 32DIM > diMNs
Biological Process
synapse organization
modulation of chemical synaptic transmission
regulation of trans-synaptic signaling
cilium movement
axonogenesis
axoneme assembly
cell-cell adhesion via plasma-membrane adhesion molecules
0
5
10
15
$-\log_{10}(FDR)$
Cellular Component
synaptic membrane
neuron to neuron synapse
postsynaptic membrane
asymmetric synapse
postsynaptic density
postsynaptic specialization
0
5
10
15
$-\log_{10}(FDR)$
Molecular Function
metal ion transmembrane transporter activity
salt transmembrane transporter activity
inorganic molecular entity transmembrane transporter activity
inorganic cation transmembrane transporter activity
monoatomic ion gated channel activity
gated channel activity
0
5
10
15
$-\log_{10}(FDR)$
d
PurMNs 32DIM > PurMNs 20DIM
Biological Process
regulation of small GTPase mediated signal transduction
cell junction assembly
small GTPase mediated signal transduction
regulation of neuron projection development
skeletal system development
axonogenesis
synapse organization
0.0
2.5
5.0
7.5
10.0
$-\log_{10}(FDR)$
Cellular Component
cell-cell junction
cell leading edge
neuron to neuron synapse
asymmetric synapse
glutamatergic synapse
postsynaptic density
0
2
4
6
$-\log_{10}(FDR)$
Molecular Function
GTPase regulator activity
nucleoside-triphosphatase regulator activity
GTPase activator activity
protein kinase activity
guanyl-nucleotide exchange factor activity
small GTPase binding
0.0
2.5
5.0
7.5
10.0
12.5
$-\log_{10}(FDR)$
Gene number
200
400
600
e
diMNs > PurMNs 32DIM
Biological Process
oxidative phosphorylation
mitochondrial translation
DNA replication
aerobic respiration
mitochondrial gene expression
DNA-templated DNA replication
proton motive force-driven mitochondrial ATP synthesis
0
5
10
15
20
$-\log_{10}(FDR)$
Cellular Component
mitochondrial protein-containing complex
mitochondrial inner membrane
organelle inner membrane
inner mitochondrial membrane protein complex
organellar ribosome
mitochondrial ribosome
0
20
40
60
$-\log_{10}(FDR)$
Molecular Function
oxidoreduction-driven active transmembrane transporter activity
structural constituent of ribosome
electron transfer activity
NADH dehydrogenase (ubiquinone) activity
NADH dehydrogenase (quinone) activity
NADH dehydrogenase activity
0.0
2.5
5.0
7.5
10.0
12.5
$-\log_{10}(FDR)$
f
PurMNs 20DIM > PurMNs 32DIM
Biological Process
oxidative phosphorylation
proton motive force-driven ATP synthesis
aerobic respiration
proton motive force-driven mitochondrial ATP synthesis
cellular respiration
ATP synthesis coupled electron transport
mitochondrial ATP synthesis coupled electron transport
0
10
20
30
$-\log_{10}(FDR)$
Cellular Component
mitochondrial protein-containing complex
mitochondrial inner membrane
organelle inner membrane
inner mitochondrial membrane protein complex
mitochondrial respirasome
respiratory chain complex
0
20
40
60
$-\log_{10}(FDR)$
Molecular Function
oxidoreduction-driven active transmembrane transporter activity
electron transfer activity
NADH dehydrogenase (ubiquinone) activity
NADH dehydrogenase (quinone) activity
NADH dehydrogenase activity
NAD(P)H dehydrogenase (quinone) activity
0
5
10
15
20
$-\log_{10}(FDR)$

**Figure S4. Contrast-specific tissue, pathway, and Gene Ontology enrichment analyses resolve the transcriptional programs underlying purified motor neuron maturation, related to Figure 3.**

(a) ARCHS4 tissue enrichment for genes elevated in purMNs 32DIM relative to diMNs. (b) KEGG pathway enrichment for the same contrast. (c-f) GO enrichment displayed by contrast, with Biological Process, Cellular Component, and Molecular Function panels shown separately for each comparison: (c) purMNs 32DIM > diMNs, (d) purMNs 32DIM > purMNs 20DIM, (e) diMNs > purMNs 32DIM, and (f) purMNs 20DIM > purMNs 32DIM. Enrichment analyses were performed on genes elevated in each indicated contrast from limma differential-expression results, using BH-adjusted $p < 0.05$ and $|\log FC| >= 1$ as the input gene threshold. GO Biological Process, Cellular Component, and Molecular Function terms were tested with clusterProfiler::enrichGO, and KEGG pathways were tested with clusterProfiler::enrichKEGG using the mapped Entrez gene universe. Bubble size denotes overlapping gene number, x-axis position denotes -log10(FDR), and darker color indicates stronger enrichment within each comparison block. Terms shown met FDR < 0.05.

**Table S1. Human iPSC lines used in this study**

**A. Cell-line and donor information**

| Cell line and parent identifier | Experimental use | Source/cohort | Cell source | Sex | Age (years) | Ethnicity | Race | Clinical/genetic information |
|---|---|---|---|---|---|---|---|---|
| CS8VTRiCTR-nxx (parent GUID: NEUXC258VTR) | diMN and purMN differentiation | Cedars-Sinai Biomanufacturing Center; Answer ALS | PBMC | Female | 28 | Not Hispanic or Latino | White | Clinically categorized control |
| CS0YX7iCTR-nxx (parent GUID: NEUVZ050YX7) | diMN and purMN differentiation | Cedars-Sinai Biomanufacturing Center; Answer ALS | PBMC | Male | 49 | Not Hispanic or Latino | White | Clinically categorized control |
| EDi022-A (RRID: CVCL_VT08) | diMN and purMN differentiation | Cedars-Sinai Biomanufacturing Center; Lothian Birth Cohort | PBMC | Male | 79 | Not Hispanic or Latino | White | Clinically categorized control |
| CSC640iCTR-n6 | SMPC differentiation and NMJ-chip co-culture | Cedars-Sinai Biomanufacturing Center; previously described by Villalba et al. | PBMC | Male | 30 | Not Hispanic or Latino | White | Clinically unaffected heterozygous carrier of a pathogenic LAMA2 variant |

**B. Available cell-line quality-control information**

| Cell line | Reprogramming method | Karyotype | Pluripotency assessment | Identity authentication | Mycoplasma |
|---|---|---|---|---|---|
| CS8VTRiCTR-nxx | Episomal plasmids | 46,XX; normal G-banded karyotype | Positive alkaline phosphatase and pluripotency-marker ICC; TaqMan hPSC Scorecard self-renewal and three-lineage assessments passed; PluriTest not reported | STR profile established as unique | Negative |
| CS0YX7iCTR-nxx | Episomal plasmids | 46,XY; normal G-banded karyotype | Positive alkaline phosphatase and pluripotency-marker ICC; PluriTest passed; TaqMan hPSC Scorecard self-renewal and three-lineage assessments passed | STR profile matched the primary donor material | Negative |
| EDi022-A | Episomal plasmids | 46,XY; normal G-banded karyotype | Positive alkaline phosphatase and pluripotency-marker ICC; PluriTest passed; Scorecard self-renewal and endoderm assessments passed, ectoderm was borderline, and mesoderm did not pass | STR profile matched the primary donor material | Negative |
| CSC640iCTR-n6 | Not reported in the available public derivative record | Not reported in the available public derivative record | Not reported in the available public derivative record | STR profile of the reported derivative matched the primary donor material | Reported derivative tested negative |

**Abbreviations:** ICC, immunocytochemistry; MN, motor neuron; NMJ, neuromuscular junction; PBMC, peripheral blood mononuclear cell; SMPC, skeletal muscle progenitor cell; STR, shorttandem repeat. Quality-control information was obtained from Cedars-Sinai Biomanufacturing Center records and was not generated as part of the present study unless otherwise indicated. NR indicates information not reported in the available records. CSC640iCTRn6 was clinically unaffected but carried one pathogenic LAMA2 allele; this line was used only to generate the SMPC component of the co-culture experiments and was not included in the motor neuron transcriptomic comparisons.

**Table S2**

| Primary antibody | Dilution | Reference | Company |
|---|---|---|---|
| MNX1/Hb9 | 1:300 | ab221884 | Abcam |
| ISL1/2 | 1:50 | 39.4D5 / AB_2314683 | DHSB |
| ChAT | 1:300 | AF3447 | R&D Systems |
| S100B | 1:500 | S2532 | Sigma-Aldrich |
| GFAP | 1:500 | Z033401-2 | Agilent Dako |
| NEFH | 1:500 | AB1989 | Sigma-Aldrich |
| MHC | 1:1000 | MAB1548 | Sigma-Aldrich |
| Alpha bungarotoxin conjugated to Alexa 647 | 1:500 | B35450 | Thermo Scientific |
